# Effects of early life adversity and adolescent basolateral amygdala activity on corticolimbic connectivity and anxiety behaviors

**DOI:** 10.1101/2024.03.26.586708

**Authors:** Caitlyn R. Cody, Emilce Artur de la Villarmois, Anabel Miguelez Fernandez, Janelle Lardizabal, Chaney McKnight, Kuei Tseng, Heather C. Brenhouse

**Affiliations:** Psychology Department, Northeastern University, Boston MA 02115; Department of Anatomy and Cell Biology, University of Illinois, Chicago IL 60612

## Abstract

Early postnatal development of corticolimbic circuitry is shaped by the environment and is vulnerable to early life challenges. Prior work has shown that early life adversity (ELA) leads to hyperinnervation of glutamatergic basolateral amygdala (BLA) projections to the prefrontal cortex (PFC) in adolescence. While hyperinnervation is associated with later-life anxiety behaviors, the physiological changes underpinning corticolimbic and behavioral impacts of ELA are not understood. We tested whether postsynaptic BLA-driven PFC activity is enhanced in ELA-exposed animals, using the maternal separation (MS) model of ELA. PFC local-field potential following BLA stimulation was facilitated in MS-exposed adolescents. Since ELA increases activity of the early-developing BLA, while the PFC exhibits protracted development, we further examined impacts of glutamatergic BLA activity during early adolescence on later-life PFC innervation and heightened anxiety. In early adolescence, MS-exposed animals exhibited decreased anxiety-like behavior, and acute adolescent BLA inhibition induced behaviors that resembled those of MS animals. To examine long-lasting impacts of adolescent BLA activity on innervation, BLA-originating axonal boutons in the PFC were quantified in late adolescence after early adolescent BLA inhibition. We further tested whether late adolescent BLA-PFC changes were associated with anxious reactivity expressed as heightened acoustic startle responses. MS rearing increased BLA-PFC innervation and threat reactivity in late adolescence, however early adolescent BLA inhibition was insufficient to prevent MS effects, suggesting that earlier BLA activity or post-synaptic receptor rearrangement in the PFC drives altered innervation. Taken together, these findings highlight both pre- and postsynaptic changes in the adolescent BLA-PFC circuit following ELA.

## 1. Introduction

Early postnatal life is a dynamic period of brain development that involves a coordinated symphony of circuit maturation. As many regions within the brain continue to develop and mature postnatally, external stimuli impact this development and regulate the degree of connectivity to other brain areas. Since subcortical limbic regions undergo substantial synapse formation and pruning during early development while higher-order areas such as the prefrontal cortex (PFC) mature throughout adolescence,^1–5^ early postnatal life and adolescence are two sensitive periods when limbic and prefrontal regions form functional connections in response to external stimuli and neuronal activity. Thus, activity within limbic regions over the course of development can affect the strength and quantity of efferent connections to cortical areas.

The basolateral amygdala (BLA) has unique integration with aversion- and reward-related circuitry and serves as the primary input nucleus of the amygdala.^6^ Monosynaptic BLA connectivity with the PFC^7–10^ increases throughout adolescence in healthy rats, facilitating the integration of emotional information with cognitive appraisals of the environment.^11^ Challenges to the early amygdala that alter local activity may therefore induce activity changes within efferent projections, which may result in changes to behavior when the circuit is recruited during adulthood.

Adolescence is a dynamic period when synapses undergo significant refinement in the frontal cortex.^12, 13^ These reported changes are thought to involve glutamatergic afferents with local remodeling of inhibitory-excitatory synaptic balance,^14, 15^ and coincide with heightened emotional regulation.^16–18^ Importantly, neuronal activity during sensitive periods directly impacts pre- and post-synaptic plasticity within the PFC.^19^ The present study therefore investigated the impacts of early life environmental inputs and adolescent BLA activity on the development of BLA-PFC circuitry and corresponding behaviors.

Theoretical frameworks regarding environmental influences on development posit that postnatal brain circuitry has evolved to “expect” an appropriate quantity and quality of caregiving stimuli in order to develop typically.^20^ Disruptions to typical development can be detrimental to the postnatal brain and can exert cascading effects on subsequent maturation.^21–24^ One such form of disruption includes early life adversity (ELA), where the species-expectant environment is violated.^25, 26^ ELA can lead to several affective consequences such as anxiety and major depressive disorder, which often emerge in adolescence, and continue through adulthood.^13, 27–34^ The maternal separation (MS) model of ELA in rodents induces corticolimbic and anxiety-related changes that resemble those reported in ELA-exposed humans. Particularly, MS induces hyperinnervation of the PFC from glutamatergic BLA projections,^18^ as well as enhanced anxiety-like behavior as measured by the acoustic startle response (ASR).^35^ These effects were seen in adolescent rodents as early as postnatal day (p)40 in male rats, and p30 in females, which corresponds to early-mid adolescence. Therefore, we hypothesized that ELA leads to pre- and post-synaptic changes to BLA-PFC connectivity via hyperactivity within the developing BLA, and that preventing heightened presynaptic BLA activity during early-mid adolescence would interrupt hyperinnervation and result in reduced anxiety behaviors.

We first determined whether MS exposure led to facilitation of BLA-evoked local-field potentials (LFP) within the PFC in early-mid adolescence. Next, to test the role that BLA activity plays in the developing innervation to the PFC, BLA activity was manipulated via an inhibitory DREADD to reduce excitatory activity during early-mid adolescence. In an exploratory analysis, an open field test was performed following acute DREADD inhibition to test whether early adolescent BLA activity regulated anxiety-like behaviors differentially after MS exposure. Histology was performed in late adolescence to determine if earlier BLA inhibition prevented the establishment of long-range glutamatergic PFC hyperinnervation. Animals were also tested in late adolescence for enhanced acoustic startle response (ASR) at baseline or following an anxiety-provoking social threat cue (playback of 22-kHz ultrasonic vocalizations), to determine whether prevention of BLA-PFC hyperinnervation was associated with prevention of enhanced ASR.

## 2. Materials and Methods

All experimental procedures were conducted in accordance with the 1996 Guide for the Care and Use of Laboratory Animals (NIH) with approval from the Institutional Animal Care and Use Committee at Northeastern University and the University of Illinois Chicago.

### 2.1 Subjects and Maternal Separation (MS)

All experimental subjects were bred in house (details in Supplemental Materials). Whole litters were randomly assigned to control (CON) or maternal separation (MS) rearing conditions. All separations lasted 4 hours each day beginning at 0900h with pups kept in a separate room from the dam. Upon completion of MS pups were reunited with their dams and littermates and left undisturbed until the next day. On p21, pups were weaned from the dam into standard rat cages with an age, sex, and condition matched partner.

### 2.2 Local Field Potential (LFP) recordings of prefrontal response to basolateral amygdala (BLA) stimulation

Early-mid adolescent (p30-45) rats were anesthetized and a concentric bipolar electrode was used to record within the prelimbic region while stimulating the BLA through a computer-controlled pulse generator (details in Supplemental Materials). Changes in the input-output curve of PFC responses were estimated by comparing the amplitude of BLA-evoked LFP at different stimulating intensities (0, 0.25, 0.5, 0.75, and 1.0 mA) using single (300µS duration) square pulses delivered every 15s in both control and MS rats.

### 2.3 Viral Injections and Chemogenetic Inhibition

In a separate cohort of male and female rats, on p26 subjects underwent a stereotaxic surgical injection of the inhibitory DREADD AAV-CAMKIIa-HM4DGi-mCherry (Addgene, Watertown, MA) to the BLA (A/P: −1.9mm, L/M: ±4.5mm, D/V: −6.75mm). Viral injections are detailed in Supplemental Materials. Rats were allowed to recover for 1 week before behavioral testing. Daily from p33-39, rats were weighed and given an intraperitoneal (i.p) injection of the DREADD agonist clozapine-N-oxide dihydrochloride (CNO, 2mg/kg, Bio-Techne Tocris, Minneapolis, MN) or vehicle saline (SAL). CNO was prepared fresh prior to the start of injections and was dissolved in sterile saline. Functional validation methods are detailed in Supplemental Materials.

### 2.4 Behavioral Assays

For each behavioral assay, rats were brought into the behavioral suite and allowed to habituate for 15 minutes.

#### 2.4.1 Open Field Test (OFT)

For early adolescent testing (p33), 30 minutes after the first CNO or SAL injection, rats were placed into the open field (100 cm x 100 cm black acrylic arena). Video monitoring of behavior was done via an overhead CCTV camera (Panasonic WV-CP500, Secaucus, NJ), and behavioral scoring was done via Noldus Ethovision 9.0 Behavioral Tracking software. The central zone of the arena was defined as the inner 30×30 cm portion of the arena (Figure 4A).

#### 2.4.2 Acoustic Startle Response (ASR)

On p54, rats were placed into a sound attenuated startle chamber (MedAssociates Inc, St. Albens, VT) and played a series of 30 105dB white noise bursts for an assessment of baseline startle. Twenty-four hours later, rats underwent a second ASR test, during which ASR was measured following exposure to a five-minute 22 kHz rat ultrasonic vocalization (USV) recording. For more details see Supplemental Materials.

### 2.5 Microscopy and Analysis

#### 2.5.1 Viral Expression

Histological procedures are detailed in Supplemental Materials. In order to quantify the amount of DREADD expression within and outside of the BLA, slices were imaged on a Zeiss Imager M.2 microscope for bolus verification. Slices with dual hemispheric expression in the BLA underwent volumetric analysis via the Cavalieri probe within StereoInvestigator. Sections containing DREADD expression were analyzed at 2.5X magnification to determine the extent of bolus infection within and spread outside of the BLA. Average BLA volume was previously measured in a subset of animals, and the volume of DREADD expression was quantified as a percent of the total BLA volume. Animals with a total bolus volume that filled <50% of the BLA were excluded from analysis.

#### 2.5.2 Axonal Innervation

The corresponding PFC tissue of all included animals was processed using immunohistochemistry (IHC) to identify mCherry expression. Sections containing the infralimbic (IL) and preimbic (PL) regions were prepared for immunohistochemistry as detailed in Falcy et al. 2020^36^ (details in Supplemental Materials). Z-stacks were analyzed for axonal bouton count and intensity.

### 2.6 Statistical Analysis

GraphPad Prism 10.0 Software was used for all data analyses and graphical representation. Sample sizes were determined by power analyses using effect sizes from prior experiments. Sex x rearing x treatment 3-way ANOVAs were conducted (with frequency as repeated measures in BLA-evoked PFC LFP analyses), and when main effects of sex or interactions with sex were observed, 2-way ANOVAs with factors of treatment and rearing were conducted for each sex with Post-hoc Tukey’s multiple comparison’s tests when appropriate. For simplicity, full statistical details are only highlighted for significant comparisons in the Results section. For all statistical details, see Supplemental Materials. Of note, three animals were removed from ASR analysis due to data file corruption.

## 3. Results

### 3.1 MS leads to facilitation of BLA-evoked local field potential in the early adult PFC

3-way ANOVA revealed a main effect of rearing (F_1,20_ = 102.0; *p* < 0.0001; partial η^2^ = 0.732) and a rearing x intensity interaction (F_4,80_ = 38.23; *p* < 0.0001; partial η^2^ = 0.563) on PFC local field potential (Figure 1A-C; n=6/group), with MS exposure facilitating BLA-evoked activity in both males (Figure 1B) and females (Figure 1C). These findings indicate that exposure to MS results in increased BLA-evoked activity within the PFC in early-mid adolescence.

**Figure 1.**
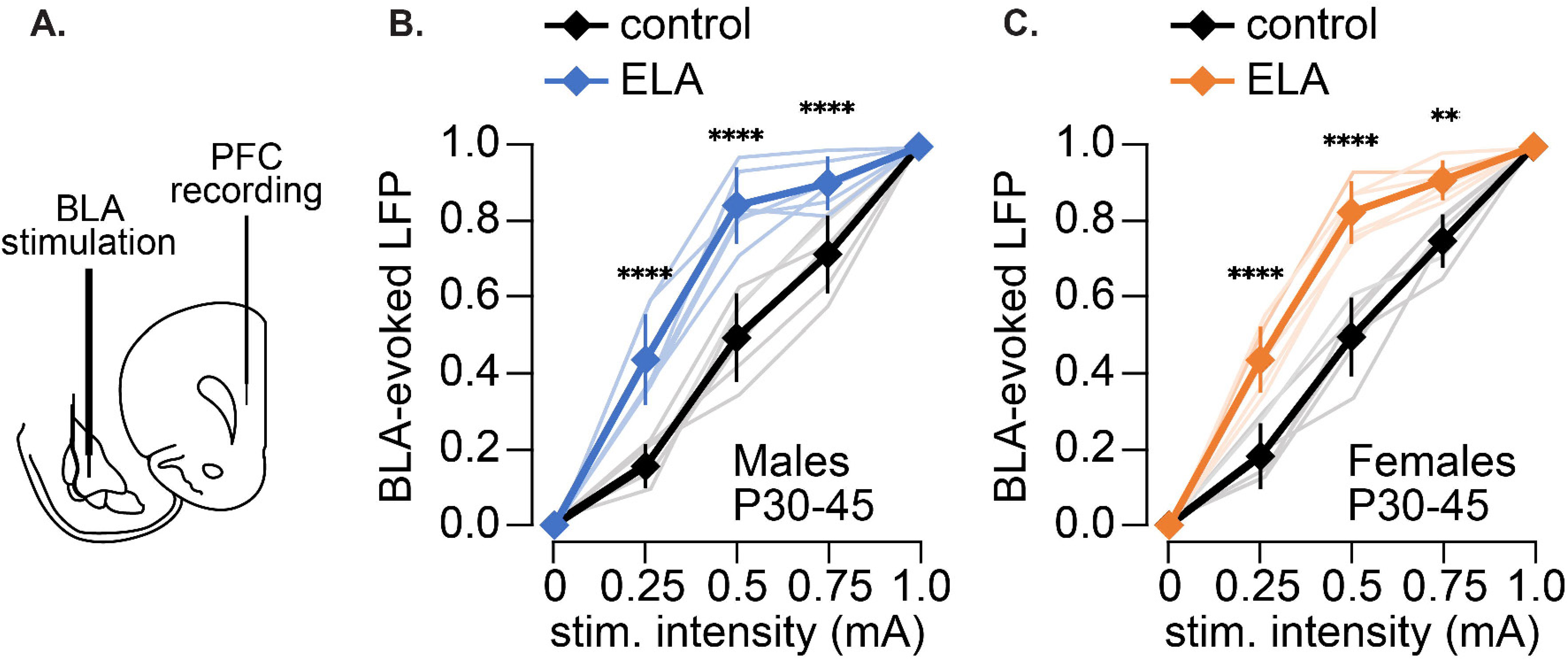
Effects of ELA on BLA-evoked PFC LFP. (**A**) Illustration of BLA stimulation and PFC recording sites. (B) Prelimbic PFC response to BLA stimulation at increasing intensities is enhanced by p30 as shown by local field potential (LFP) recordings in males (****p<0.0001, **p<0.01 difference between control and MS animals, after correction for multiple comparisons; N=6/group). (**C**) The prelimbic PFC LFP response to BLA stimulation is also enhanced by p30 in females (****p<0.0001, **p<0.01 difference between control and MS animals, after correction for multiple comparisons; N=6/group).

### 3.2 BLA inhibition during mid-adolescence does not prevent increased BLA-PFC connectivity after MS

Figure 2A summarizes the DREADD strategy used to inhibit the BLA. Ex-vivo recordings of BLA pyramidal neurons show the effect of the inhibitory DREADD at 8-9 days post viral injection (Figure 2B-D).

**Figure 2.**
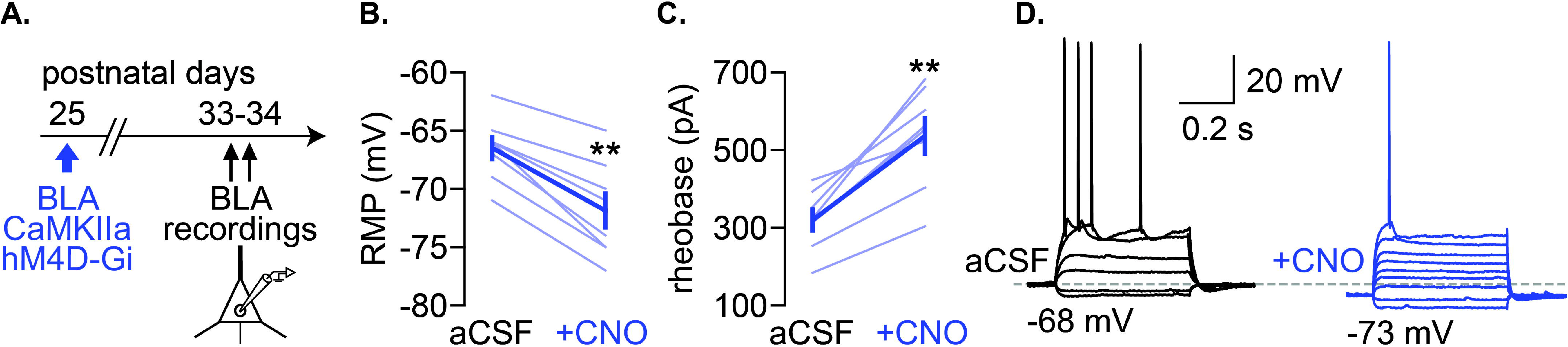
DREADD validation. (**A**) Timeline of DREADD delivery and ex-vivo recordings of BLA pyramidal neurons. (**B**) Bath application of CNO (10µM, 10 min) markedly hyperpolarized the resting membrane potential of all pyramidal neuron tested (**p<0.01, unpaired t-test, t=3.114, df=14). (**C**) Accordingly, a significant increase in the rheobase (minimal somatic current (pA) to elicit an action potential) was observed all neurons following bath application of CNO (**p<0.01, unpaired t-test, t=4.124, df=14). (**D**) Example traces of a BLA pyramidal neuron showing the inhibitory effect of CNO as revealed by the reduced current evoked action potential and hyperpolarized resting membrane potential.

A subset of rats (n=6-8) were sacrificed following behavioral testing (p56-60) to quantify BLA innervation to the PFC via axonal bouton count and fluorescent intensity. Three-way ANOVA revealed a sex x rearing interaction with a moderate effect size on bouton count in the PL (*F*_1,57_ = 3.245; p=0.0770; partial η^2^ = 0.054) and the IL (*F*_1,57_ = 3.617; p=0.0623; partial η^2^ = 0.06). Due to the presence of a moderate effect size for the sex x rearing interaction, each sex was analyzed separately, with two-way ANOVAs assessing effects of rearing condition and treatment (CNO or SAL). In males, MS increased PL bouton count (*F*_1,25_ = 4.661; p=0.0407; partial η^2^ = 0.157; Figure 3E) and IL bouton count (*F*_1,25_ = 5.098; p=0.0329; partial η^2^ = 0.169; Figure 3I). However, no effects were observed in females.

**Figure 3.**
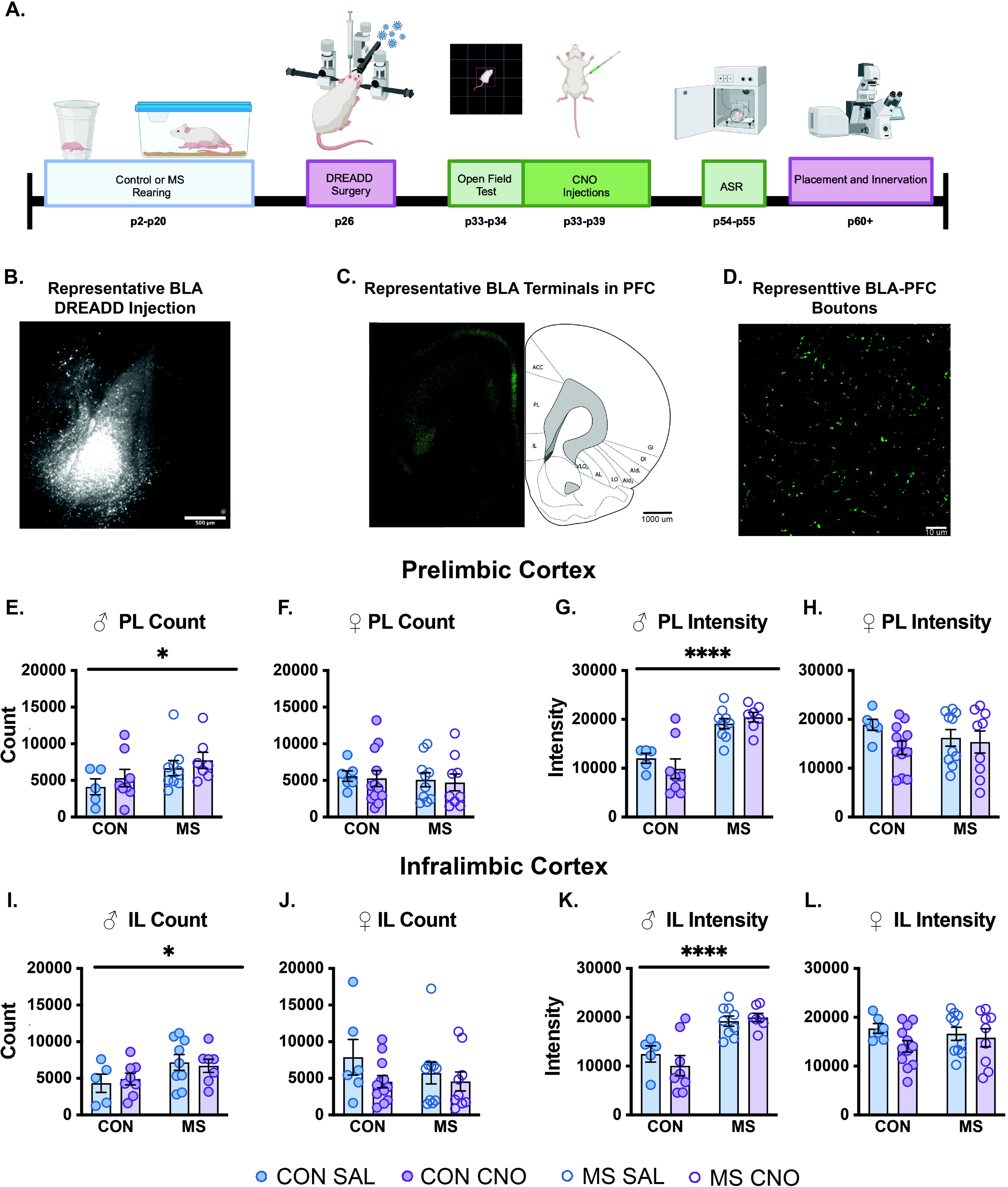
Effects of ELA with or without BLA inactivation during early-mid adolescence on late adolescent PFC innervation. **(A)** Timeline of DREADD experiment. **(B)** Representative DREADD injections in the BLA **(C-D)** BLA-PFC terminals in the PFC. **(E)** Male PL bouton count. **(F)** Female PL bouton count. **(G)** Male PL bouton intensity. **(H)** Female PL bouton intensity. **(I)** Male IL bouton count. **(J)** Female IL bouton count. (**K)** Male IL bouton intensity. **(K)** Female IL bouton intensity. Means ± SEM with individual data points are shown. (*p < 0.05 ****p<0.0001 rearing effect). n=5-12/group.

Three-way ANOVA also revealed a main effect of rearing on bouton intensity in the PL (*F*_1,57_ = 10.87; p=0.0017; partial η^2^ = 0.160) and IL (*F*_1,57_ = 15.58; p=0.0002; partial η^2^ = 0.215), as well as a sex x rearing interaction in bouton intensity within the PL (*F*_1,57_ = 15.00; p=0.0003; partial η^2^ = 0.208) and IL (*F*_1,57_ = 13.19; p=0.0006; partial η^2^ = 0.160). Due to the presence of a sex x rearing interaction, each sex was analyzed separately, revealing another male-specific main effect of rearing on PL bouton intensity (*F*_1,25_ = 36.09; p=<0.0001; partial η^2^ = 0.591, Figure 3G), and IL bouton intensity (*F*_1,25_ = 29.53; p=<0.0001; partial η^2^ = 0.541; Figure 3K).

Treatment with CNO had no effects on either measure (count or fluorescent intensity). Results of all statistical comparisons are summarized in Supplementary Tables 5-7. These findings indicate that MS rearing induces hyperinnervation of the PFC from the BLA in late adolescent male animals only, and that BLA inhibition during early-mid adolescence is insufficient to reverse these changes.

### 3.3 Exposure to MS and acute BLA inhibition alters adolescent behavior in a sex dependent manner

Rats with inhibitory DREADD expression in the BLA were given an i.p injection of CNO (2 mg/kg) or SAL 30 minutes before the open field test on p33. Animals (n=10-12 per condition) were assessed for time spent, latency to enter, and number of entries into the center zone of the arena (Figure 4D-K).

**Figure 4.**
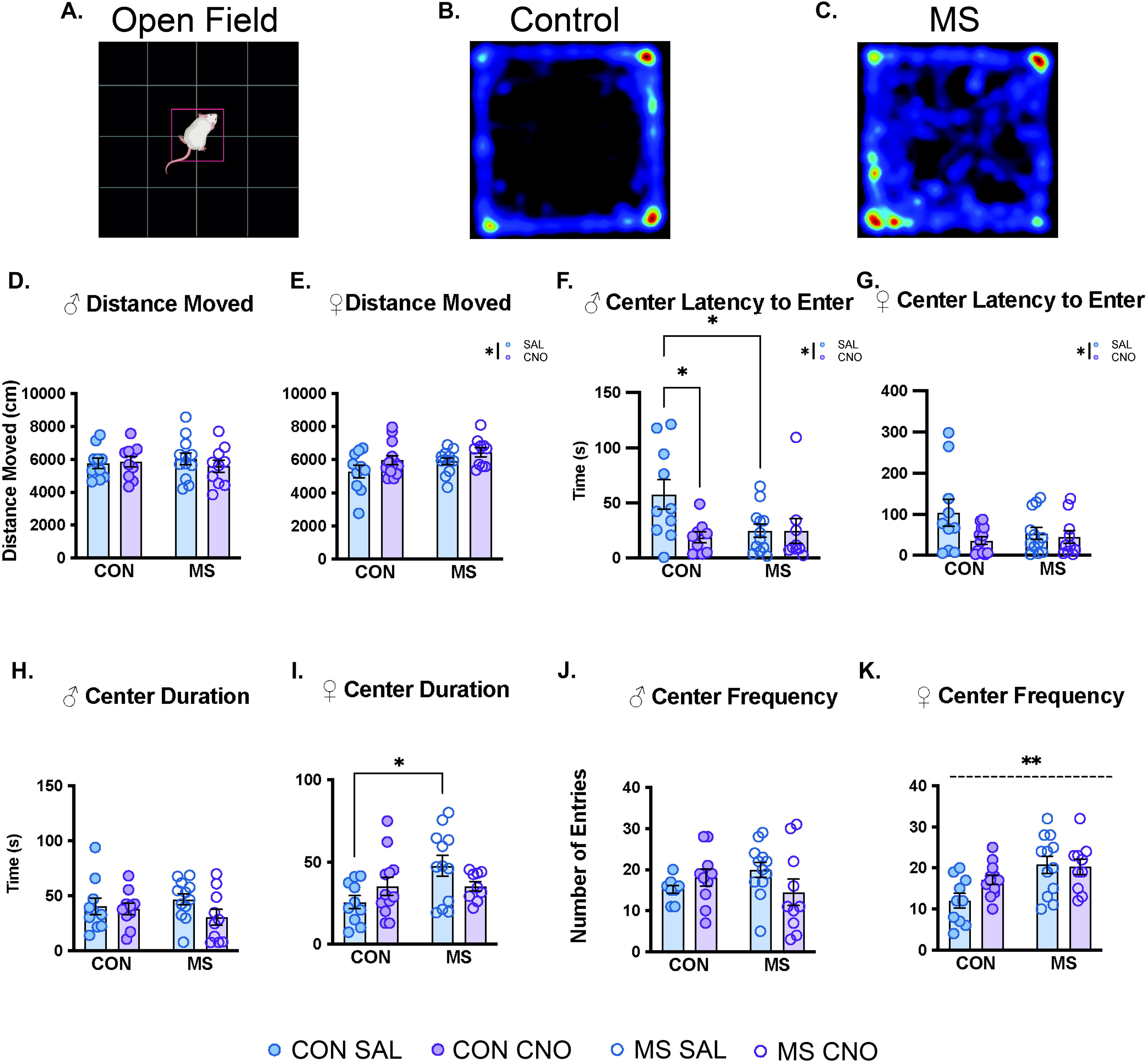
Effects of ELA with or without acute BLA inactivation on open field behavior during early adolescence. **(A)** Schematic of open field arena. **(B-C)** Representative heat maps depicting control and MS animal movement within the open field arena. OFT results for male **(D)** Distance Moved **(H)** Center Duration **(F)** Center latency to enter **(J)** Center frequency and female **(E)** Distance Moved **(I)** Center Duration **(G)** Center latency to enter **(K)** Center frequency. Means ± SEM with individual data points are shown. (*p < 0.05; **p<0.005; dashed line denotes main effect of rearing). n=10-13/group.

A three-way ANOVA revealed a moderate effect size sex x treatment interaction (*F*_1,79_ = 3.061; p=0.081; partial η^2^ = 0.037) on distance moved. Two-way ANOVAs within each sex assessed effects of rearing condition and treatment (CNO or SAL). No effects were seen in male animals (Figure 4D). In female animals, a main effect of treatment (F_1,41_= 4.526; p=0.0394; partial η^2^ = 0.099) was found indicating that females treated with CNO had increased locomotion (Figure 4E).

Three-way ANOVAs revealed main effects of sex (*F*_1,79_ = 7.019; p=0.0098; partial η^2^ = 0.084) and treatment (*F*_1,79_ = 7.148; p=0.0092; partial η^2^ = 0.085) for the latency to enter the center. Analyses also revealed a rearing x treatment interaction (*F*_1,79_ = 5.098; p=0.0268; partial η^2^ = 0.062). Two-way ANOVAs assessed effects within each sex of rearing condition and treatment. In male animals, a main effect of treatment (*F*_1,38_ = 4.475; p=0.0414; partial η^2^ = 0.11) and a rearing x treatment interaction (*F*_1,38_ = 4.235; p=0.0469; partial η^2^ = 0.105) were found for the latency to enter the center (Figure 4F). Post-hoc analyses revealed differences between control SAL and control CNO treated animals (p=0.0129) as well as control SAL and MS SAL animals (p=0.0258) indicating that control animals treated with CNO as well as MS animals treated with SAL exhibited a decrease in the latency to enter the center compared to control SAL animals. In female animals, a significant main effect of treatment (*F*_1,41_ = 4.185; p=0.0472; partial η^2^ = 0.093) was found in the latency to enter the center (Figure 4G), with CNO treated animals showing a decreased latency to enter the center.

Three-way ANOVAs revealed a rearing x treatment interaction for center duration (*F*_1,79_ = 4.912; p=0.0296; partial η^2^ = 0.059). Two-way ANOVAs further assessed effects of rearing condition and treatment within each sex. No effects were seen in male animals (Figure 4H). In female animals, analyses revealed a main effect of rearing (*F*_1,41_ = 4.392; p=0.0427; partial η^2^ = 0.101) and a rearing x treatment interaction (*F*_1,41_ = 4.412; p=0.0422; partial η^2^ = 0.101) on center duration (Figure 4I). Post-hoc analyses revealed differences between control and MS SAL treated females (p=0.0045) such that MS animals spent more time in the center compared to controls, and that this was not mediated by CNO treatment.

A three-way ANOVA revealed a main effect of rearing (*F*_1,79_ = 5.353; p=0.0234; partial η^2^ = 0.065) and a rearing x treatment interaction (*F*_1,79_ = 6.176; p=0.0151; partial η^2^ = 0.074) for center frequency. Two-way ANOVAs within each sex assessed effects of rearing condition and treatment. In male animals, a moderate effect sized rearing x treatment interaction was observed (*F*_1,38_ = 3.561; p=0.0677; partial η^2^ = 0.087; Figure 4J). Female animals displayed a significant main effect of rearing (*F*_1,41_ = 11.24; p=0.0018; partial η^2^ = 0.219; Figure 4K), such that MS females entered the center more than control-reared females, mirroring the increased duration. These findings indicate that MS rearing results in decreased anxiety-like behavior in the OFT, and that BLA inhibition in early adolescence results in an MS-like phenotype in control animals, while having no effect on MS animals.

### 3.4 Early adolescent BLA inhibition does not prevent ASR changes in MS-exposed late adolescence

On p54 (late adolescence), rats were tested for baseline startle levels to 30 white noise bursts. Latency to startle, peak startle time, and peak startle value were assessed with three-way ANOVAs. Previously established differences between males and females were observed and are likely related to animal size; main effects of sex were observed on baseline latency to startle (*F*_1,76_ = 7.821; p=0.0065; partial η^2^= 0.093), and baseline peak value (*F*_1,76_ = 5.598; p=0.0205; partial η^2^= 0.069). No effects of rearing or treatment were observed on baseline ASR in late adolescence (Figure 5A-C). On p55, rats were presented with a five-minute playback of a 22 kHz USV recording and then immediately tested for startle levels to 30 white noise bursts. Latency to startle, peak startle time, and peak startle value were assessed with three-way ANOVAs. Main effects of rearing were found for latency to startle (*F*_1,76_ = 4.855; p=0.0306; partial η^2^= 0.060; Figure 5D), and peak time (*F*_1,76_ = 5.210; p=0.0252; partial η^2^ = 0.064; Figure 5E), with MS-exposed animals startling more quickly than controls following USV playback. Main effects of sex were found for latency to startle on Day 2 (*F*_1,76_ = 4.082; p=0.047; partial η^2^= 0.051) as well as peak value on Day 2 (*F*_1,76_ = 10.54; p=0.0017; partial η^2^= 0.122). No effects of treatment were observed on USV mediated ASR. Rats were also evaluated for the percent change from baseline following USV presentation. A main effect of rearing was again found in latency to startle (*F*_1,76_ = 4.081; p=0.0469; partial η^2^ = 0.051; Figure 5G) and peak time (*F*_1,76_ = 5.195; p=0.0255; partial η^2^ = 0.064; Figure 5H), indicating that MS animals startled quicker than control animals following USV playback. A main effect of rearing was also observed for percent change in peak value (*F*_1,76_ = 4.856; p=0.0306; partial η ^2^= 0.060) No main effects of treatment with CNO were found in any analyses, and no interactions of rearing, sex, or treatment were present, therefore further analyses were not conducted. These findings indicate that BLA inhibition via CNO in early-mid adolescence had no overall effect on ASR in late adolescence.

**Figure 5.**
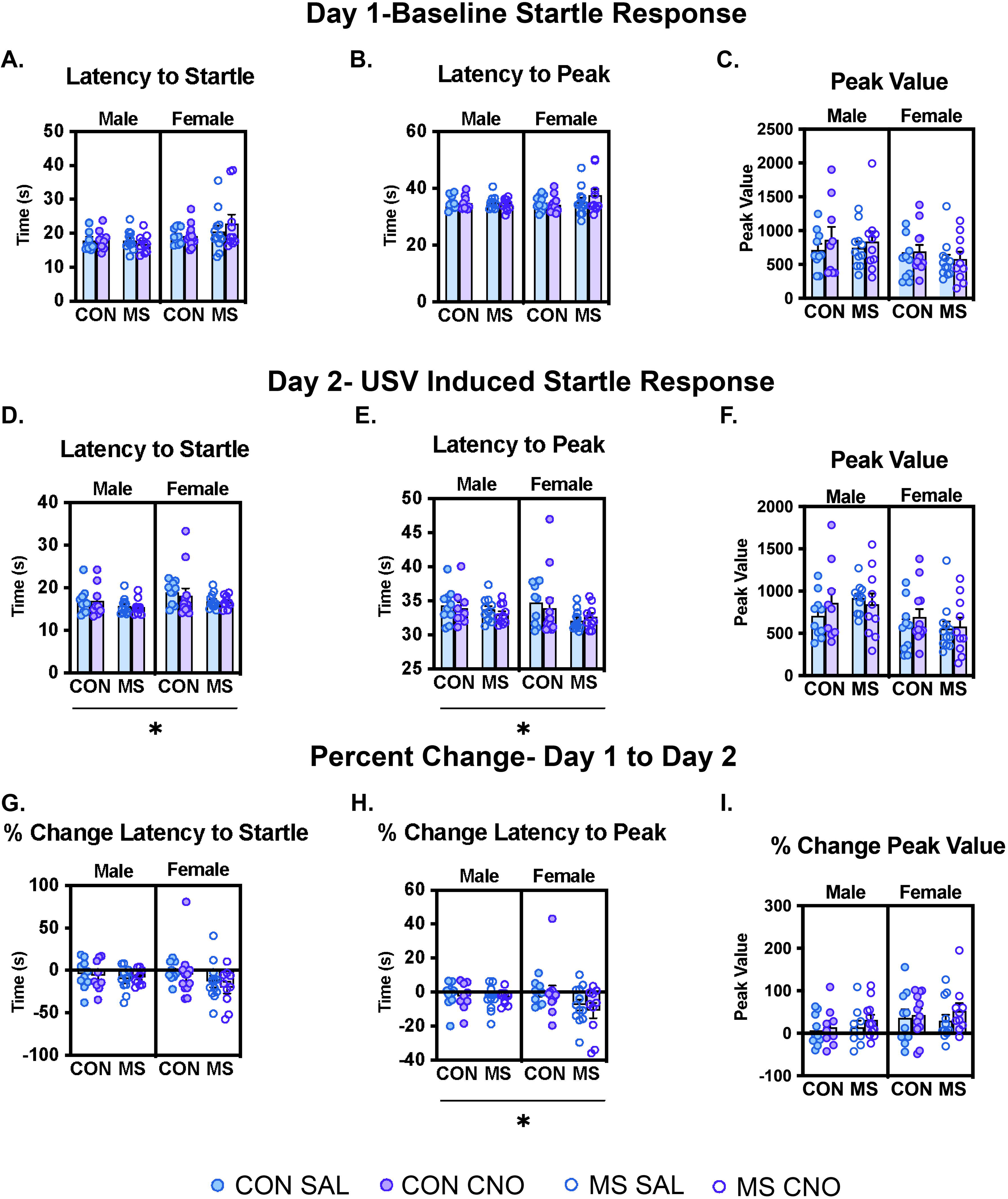
Effects of ELA with or without BLA inactivation during early-mid adolescence on late adolescent acoustic startle. (**A**) Day 1 latency to startle. (**B**) Day 1 latency to peak startle value. **(C)** Day 1 peak startle value. **(D)** Day 2 latency to startle. (**E**) Day 2 latency to peak startle value. **(F)** Day 2 peak startle value. **(G)** Percent change latency to startle. (**H**) Percent change latency to peak startle value. **(C)** Percent change peak startle value. Means ± SEM with individual data points are shown. (*p < 0.05 rearing effect). n=9-12/group.

## 4. Discussion

The current study identifies dissociable pre- and post-synaptic contributions to sex- and age-specific effects of ELA on corticolimbic connectivity and behavior. We sought to investigate if an adverse rearing environment altered BLA-PFC connectivity via aberrantly high adolescent BLA activity, and if these changes drove anxiety-like behavioral responses in early and late adolescence. We observed that MS rearing yielded postsynaptic facilitation of PFC responsivity to BLA activation by mid-adolescence. We hypothesized that presynaptic inhibition of the BLA during this period would prevent MS-driven hyperinnervation from the BLA to the PFC, as well as MS-driven changes to anxiety-like behaviors.

As previously observed by quantifying BLA axon terminals,^18^ here, MS resulted in hyperinnervation of the PFC as assessed by a more functionally relevant measure, namely the number of axonal boutons. However, this increase was not attenuated by BLA inhibition from p33-p39, suggesting that sustained adolescent activity of the BLA does not regulate the development of presynaptic connectivity with the PFC. Additionally, no increases in innervation were seen in females, indicating that by late adolescence, any previously observed increases in BLA-PFC connectivity may have pruned back to control levels. Both male and female rats exhibit significant pruning within the PFC between adolescence and adulthood, however females display greater losses than males.^37^ The age at which peak synaptic pruning of the PFC occurs in males and females remains unknown,^38^ which allows for the possibility that inhibition of BLA activity in early-mid adolescence had differential effects on BLA-PFC innervation across sexes. It is therefore possible that the period of accelerated BLA-PFC innervation induced by MS rearing occurred at an earlier timepoint in females, and thus by the time histology was performed (p56), both control and MS females had reached adult innervation levels. Additionally, previous work from our group showed increases in female BLA-PFC innervation that were already apparent by p28; therefore, if BLA activity does provoke innervation to the PFC in females, inhibition from p33-39 may have been too late to affect any MS-evoked changes. Thus, it will be important to repeat this study with an earlier intervention time point as well as earlier ages of assessment to capture any sex-specific effects of juvenile BLA inhibition on adolescent innervation.

Since inhibiting presynaptic activity was not sufficient to reverse long-range hyperinnervation, it is possible that postsynaptic plasticity within the PFC precedes and drives these connectivity changes. For example, MS results in heightened expression of GluN2A-containing NMDA receptors within the PL^39^ as well as reduced GluA2-positive AMPA receptors.^40^ While the PFC displays late-maturing connectivity with regions such as the BLA,^11, 18^ the postnatal environment is capable of directly impacting PFC activity and receptivity,^41, 42^ and the nature of glutamate receptivity can influence development of inputs.^43^ Importantly, post synaptic NMDAR increases induced by MS have been shown to correlate with anxiety-like behaviors,^44^ and thus in concert with our findings here showing a lack of structural changes following presynaptic inhibition, we posit that postsynaptic mechanisms may be more critical for ELA induced outcomes in the PFC.

We observed anxiety-like ASR in late adolescence such that MS animals responded to anxiogenic USV playback with elevated threat responsivity. Contrastingly, MS-exposed animals tested in early adolescence exhibited faster center entry (males) or increased center time (females), suggesting a decrease in anxiety-like behavior. Our findings are largely consistent with other assessments of MS-reared animals in early adolescence,^45–47^ where by late adolescence, MS-reared animals display reduced center time.^48^ Importantly, the open field task has also been used as a measure of risk-seeking,^49^ and ELA is known to increase risk-seeking and impulsivity in adolescence.^50–59^ It is possible therefore that the anxiolytic behaviors seen during the OFT in early adolescence may be attributable to increases in risk-seeking behaviors, and that this behavioral phenotype shifts as animals continue to mature. Notably, acute BLA inhibition in adolescent controls resulted in open field behavior that mirrored MS-exposed animals, such that control animals treated with CNO had shorter latencies to enter the center of the open field arena. Further work will tease apart the discrete mechanisms linking MS to risk-seeking, and how developing BLA activity differentially drives these behaviors during adolescence.

MS does not appear to alter later-life ASR via hyperactivity of the developing BLA, as early-mid adolescent BLA inhibition did not prevent MS-induced changes. The amygdala regulates anxiety-related enhancement of ASR reflex, with the BLA particularly sensitive to ambiguous or distal threats from the environment.^60, 61^ Here we tested the impacts of MS and early BLA manipulation on potentiation of ASR by a distal social threat, namely playback of 22kHz USV.^57^ Previous work has shown enhanced BLA activity following presentation of a 22 kHz USV,^62^ indicating a role of the BLA in the processing of this fearful stimuli. Control-reared animals display a potentiated ASR following USV playback, while MS alters this potentiated response.^35^ Here we observed that while MS-exposed animals exhibited increased potentiation of the ASR, early-mid adolescent BLA inhibition was not sufficient to rescue the heightened startle response induced by MS.

These findings should be considered with the limitation that surgery and chronic injections, which are stressful experiences in and of themselves, were conducted during juvenility and early adolescence. This may be considered a second hit of stress later following MS exposure, which can alter typical stress responses.^63^ We are also unable to causally link altered BLA-PFC innervation or functional connectivity with ASR. Taken together, we observed that postnatal ELA impacts development of both pre- and post-synaptic BLA-PFC connectivity as well as anxiety-related behaviors across adolescence. However, the insufficiency of pre-synaptic BLA inhibition to alter later BLA-PFC connectivity or anxiety-related behavior highlights the importance of investigating early post-synaptic changes in the PFC that may be driving MS-induced corticolimbic functional connectivity changes.

## Acknowledgments

The authors would like to thank the Institute for Chemical Imaging of Living Systems at Northeastern University for consultation and imaging support.

## Author Contributions

Heather C. Brenhouse and Kuei Y. Tseng and designed the study. Caitlyn R. Cody performed and analyzed the behavioral and axonal innervation data under the supervision of Heather C. Brenhouse. Janelle Lardizabal and Chaney McKnight assisted Caitlyn R. Cody with behavioral experiments, histology, and data collection. Emilce Artur de la Villarmois and Anabel Miguelez Fernandez performed all electrophysical experiments and data analysis under the supervision of Kuei Y. Tseng. Caitlyn R. Cody, Heather C. Brenhouse, and Kuei Y. Tseng wrote the manuscript.

Caitlyn R. Cody, Emilce Artur de la Villarmois, and Kuei Y. Tseng prepared the figures.

## Funding

Supported by NIH Grant R01MH127850-01 (KYT and HCB)

## Competing Interests

The authors have nothing to disclose

## Notes

### Competing Interest Statement

The authors have declared no competing interest.

### Summary of Updates

Author affiliations updated, supplemental files updated, figures updated, manuscript updated to add more detail.

