## Supplemental Materials for "Effects of early life adversity and adolescent basolateral amygdala activity on corticolimbic connectivity and anxiety behaviors"

### Supplemental Methods

*Detailed Subjects and Maternal Separation (MS)*

All rats were housed under standard laboratory conditions in a 12h light/dark cycle (lights on at 0700h), in a temperature and humidity-controlled room with access to food and water ad libitum. All experimental subjects were bred in house. Male and female rats were co-housed, and pregnancy was assessed daily. Males were removed once pregnancy was confirmed. Dams gave birth on p0. Litters were called on p1 to 12 pups (6 male) when possible, toe clipped on p5 for animal identification, and weaned on p21. Whole litters were randomly assigned to control (CON) or maternal separation (MS) rearing conditions. Pups in control litters were left undisturbed except for the husbandry procedures described above and regular weight recording and handling twice per week. Pups in the MS condition were separated from their dams and littermates in individual cups with home cage pine shavings within a circulating water bath thermo-regulated at 37C from P2-P10. At P11-20, when body temperature can be self-regulated, pups were separated into individual mouse cages. All separations lasted for a total of 4 hours each day beginning at 0900h and pups were kept in a separate room from the dam. Of note, animals tested for BLA-evoked PFC LFP were exposed to MS from P11-20 only. Upon completions pups were reunited with their dams and littermates and left undisturbed until the next day. On p21, pups were weaned from the dam into standard rat cages with an age, sex, and condition matched partner.

*Detailed Local Field Potential (LFP) recordings of prefrontal response to basolateral amygdala (BLA) stimulation*

Early-mid adolescent (p30-45) rats were anesthetized and a concentric bipolar electrode was used to record within the prelimbic region while stimulating the BLA through a computer-controlled pulse generator.  with 8% chloral hydrate (400 mg/kg, i.p.), placed in a stereotaxic apparatus (ASI Instruments, MI), and maintained at 37–38 °C using a Physitemp TCAT-2LV Controller (Physitemp Instruments, NJ), as previously described (Thomases et al., 2013; Cass et al., 2013). All recordings were conducted in the prelimbic region using a concentric bipolar electrode (SNE-100 × 50 mm; Rhodes Medical Instruments, CA) while stimulating the BLA (concentric bipolar electrode NE-100 × 50 mm; Rhodes Medical Instruments, CA) through a computer-controlled pulse generator (Master-8 AMPI, Jerusalem, Israel). Changes in the input-output curve of PFC responses were estimated by comparing the amplitude of BLA-evoked LFP at different stimulating intensities (0, 0.25, 0.5, 0.75, and 1.0 mA) using single (300$\mu$S duration) square pulses delivered every 15s in both control and MS rats.

*Functional Validation of DREADD in the BLA*

All brain slicing procedures were conducted as previously described (Flores-Barrera et al., 2017; 2020). Specifically, coronal brain slices (350$\mu$m-thick) containing the BLA were obtained within 6-7 days post-DREADD delivery to validate the functional expression of the hM4D-Gi receptor on amygdalar pyramidal neurons. All recordings were conducted at 33–35 °C in artificial cerebrospinal fluid (aCSF) containing (in mM): 122.5 NaCl, 3.5

KCl, 25 $NaHCO_3$, 1.0 $NaH_2PO_4$, 2.5 $CaCl_2$, 1.0 $MgCl_2$, 20 glucose, 1.0 ascorbic acid (pH 7.40–7.43, 295–305 mOsm) under constant oxygenation with 95 % $O_2$–5 % $CO_2$. The patch electrode was pulled from borosilicate glass (8-10 MΩ) and filled with a potassium-based internal solution containing 0.125 % Neurobiotin and (in mM): 115 K-gluconate, 10 HEPES, 20 KCl, 2.0 $MgCl_2$, 2.0 Mg-ATP, 2.0 $Na_2$-ATP, 0.3 $Na_2$-GTP (pH 7.23–7.28, 280–282 mOsm). After 5 min of obtaining the whole-cell configuration, a set of somatic current pulses was applied in current-clamp mode to measure changes in neuronal excitability (input resistance, rheobase, latency to the first spike) prior and after bath application of the DREADD ligand CNO (10$\mu$M, 10 min).

*Viral Injections for Chemogenetic Inhibition*

In a separate cohort of male and female rats, on p26 subjects underwent a stereotaxic surgical injection of the inhibitory DREADD AAV-CAMKIIa-HM4DGi-mCherry (Addgene, Watertown, MA) to the BLA (A/P: -1.9mm, L/M: ±4.5mm, D/V: -6.75mm). Rats were anesthetized with 5% isoflurane mixed with oxygen, placed in a stereotaxic apparatus (Kopf, Tujunga, CA) and maintained at 37-38°C using a heating pad. Constant level of isoflurane (2–3%) was delivered during the surgical procedure and the level of anesthesia was monitored closely by breathing rate and the pedal reflex. After exposing the skull, two small (1 mm) holes were drilled above the M/L and A/P coordinates of the BLA. A Hamilton syringe was slowly lowered to the D/V coordinates and 200 nL of virus was injected at a rate of 50 nL/min via a Harvard Apparatus Pump 11 Elite Microinfusion pump for a total of 4 minutes. The Hamilton was left at the surgical site for 1 additional minute, for a total of 5 minutes/hemisphere, before slowly being withdrawn. The surgical incision was closed with sterile sutures and treated with triple antibiotic ointment. Rats were placed back into the home cage following the return of movement and reflexes.

*Histology*

Following all behavioral testing, rats were deeply anesthetized and transcardially perfused with ice cold saline and 4% paraformaldehyde (PFA) solutions. Brains were extracted and preserved in 4% PFA for 24 hours in individual scintillation vials, followed by cryoprotection in 30% sucrose solution until brains reached optimal density. Brains were then cryosectioned on a Leica 6800 cryostat in 100um sections to isolate the PFC and BLA regions within 1 week of cryoprotection. Slices were placed in well plates with PBS until mounting and staining. Sections were collected in 6 well increments (600um). BLA slices were mounted on Fisherbrand ColorFrost Plus slides.

*Axonal Innervation*

Sections were stained with a rabbit anti mCherry primary antibody (1:2500, Abcam ab167453) and a Alexafluor 488 Goat anti Rabbit secondary antibody (1:1000, Jackson ImmunoResearch Laboratories). Three 100um slices of the PFC were imaged on a LM800 Zeiss confocal microscope. Four Z-stacks were taken in three representative PFC slices, totaling 12 z-stacks per animal. Z-Stacks were taken in .5um steps between image planes with a 63X/ 1.4 oil objective with a pinhole size of 1 Airy unit. Images were acquired with a voxel size of 0.09 um x 0.09 um x 0.5. Image stacks were 3-D reconstructed using Arivis Vision4D software (Zeiss) and analyzed for bouton detection via the BlobFinder pipeline. Settings were determined using representative stacks from all groups and optimized for bouton detection with minimal noise.

*Acoustic Startle Response Testing*

Acoustic startle response (ASR) testing protocols were adapted from and can be found in Granata et al., 2022. Male and female rats with AAV-CAMKIIa-HM4DGi-mCherry BLA injections reared under control or MS conditions and treated with saline or CNO in early-mid adolescence were tested on p54-55 in a 2-day paradigm. Because the rats' weights differ between sexes, the startle boxes were calibrated to accommodate the weight of each rat being tested.

All ASR testing was performed with the acoustic startle hardware and software package from Med Associates (Med Associates product number: MED-ASR-PRO1). The startle cabinets were equipped with sound attenuating foam on all walls and doors. Each cabinet held a grid rod animal holder on a startle platform containing the load cell. The load cell and load cell amplifier were used to convert force on the platform to a voltage representing the startle response. Speakers for delivering white noise background and startle noise bursts were positioned 1" behind the animal holder. The grids rods on the back of the animal holder provide ventilation and do not interfere with sounds. Four different chambers were counterbalanced between sexes and rearing groups.

The purpose of Day 1 was to establish a baseline startle response for each animal. On Day 1, rats were transported to the testing room (~50 lux) and left to acclimate for 15 minutes. After acclimation, each rat was placed into the animal holder in the unlit startle cabinet. The experiment began with 5 min of white background noise to acclimate the animal to the startle cabinet. Then, 30 stimulus tones were presented at 30-s intervals. Each stimulus was a 50 ms white noise tone of 105 dB with a 3 ms rise/fall time. The background noise level was set to 70 dB throughout the experiment. At the end of 30 trials, rats were removed from the boxes and returned to their home cages. Boxes were cleaned with 50% EtOH between runs.

The purpose of Day 2 was to determine the rats' response to startle stimulus tones after being exposed to a 22 KHz USV, a social cue signifying potential threat. Twenty-four hours after the Day 1 test, rats were transported to the testing room and left to acclimate for 15 min. Similar to Day 1, rats were placed in the animal holders, and the experiment began with a 5-min acclimation period. Then, rats were presented with a 5-min 22 KHz USV recording previously obtained from an adult male rat exposed to cat urine. The USV was recorded using a condenser ultrasonic microphone (Avisoft-Bioacoustics CM16/CMPA; frequency range 2–200 KHz) and played back using a D/A converter/power amplifier and ultrasonic speaker (Avisoft-Bioacoustics USG Player 416H, Vifa speakers) situated inside the startle cabinet 1" behind the animal holder. After the USV playback, rats were presented with 30 stimulus tones at 30-s intervals. Stimulus tones were 50 ms in duration, 105 dB in volume, with a 3 ms rise/fall time. At the end of Day 2, rats were removed from the boxes and returned to their home cages.

Onset measured represent the average across all trials on Day 1 or Day 2, or the percent change from Day 1 to Day 2, for each subject. For baseline measures, we report latency to onset (ms), latency to peak (ms), and peak startle value (arbitrary units), described in Supplementary Table 1.

| Dependent Variable | Definition |
| --- | --- |
| Latency to Onset (ms) | Time between white noise burst onset and response onset |
| Latency to Peak (ms) | Time between white noise burst onset and response peak |
| Peak Value (arbitrary units) | Greatest startle amplitude in the response window |

Supplementary Table 1. Description of outcome measured from the acoustic startle test.

| Dependent Variable | Comparison | F | p | Effect Size ($\eta_p^2$) | Post-hoc |
|---|---|---|---|---|---|
| BLA-evoked PFC LFP | **Frequency** | **$F_{(2.492, 48.85)}$=1050** | **<0.0001** | **0.98131427** | **-** |
| | Sex | $F_{(1,20)}$=.1169 | 0.7360 | 0.00280586 | - |
| | **Rearing** | **$F_{(1,20)}$=102** | **<0.0001** | **0.71062342** | **-** |
| | Frequency x Sex | $F_{(4,80)}$= 0.2241 | 0.9242 | 0.01108054 | - |
| | **Frequency x Rearing** | **$F_{(4,80)}$= 38.23** | **<0.0001** | **0.65656333** | **-** |
| | Sex x Rearing | $F_{(1,20)}$=.2241 | 0.6411 | 0.00536369 | - |
| | Frequency x Sex x Rearing | $F_{(4,80)}$= 0.07551 | 0.9895 | 0.00376097 | - |

Supplementary Table 2. Results of 3-way ANOVAs for frequency, sex, and rearing effects on BLA-evoked LFP in the PFC. Significant effects are highlighted in dark grey and bolded, trending effects are highlighted in light grey.

| Dependent Variable | Comparison | F | p | Effect Size ($\eta_p^2$) | Post-hoc |
|---|---|---|---|---|---|
| Male BLA-evoked PFC LFP | **Frequency** | **375** | **<0.0001** | **0.96774993** | **-** |
| | Rearing | 77.05 | **<0.0001** | **0.60647308** | **-** |
| | Interaction | 14.72 | **<0.0001** | **0.5407695** | **.25 Hz: CON vs MS p=<0.0001 .5 Hz: CON vs MS p=<0.0001 .75 Hz: CON vs MS p=<0.0001** |

Supplementary Table 3. Results of 2-way ANOVAs for frequency and rearing effects on BLA-evoked LFP in the PFC in male animals. Significant effects are highlighted in dark grey and bolded, trending effects are highlighted in light grey.

| Dependent Variable | Comparison | F | p | Effect Size ($\eta_p^2$) | Post-hoc |
|---|---|---|---|---|---|
| Female BLA-evoked PFC LFP | **Frequency** | **541** | **<0.0001** | **0.97741831** | **-** |
| | Rearing | 91.71 | **<0.0001** | **0.64719626** | **-** |
| | Interaction | 18.24 | **<0.0001** | **0.59335727** | **.25 Hz: CON vs MS p=<0.0001 .5 Hz: CON vs MS p=<0.0001 .75 Hz: CON vs MS p=0.0011** |

Supplementary Table 4.  Results of 2-way ANOVAs for frequency and rearing effects on BLA-evoked LFP in the PFC in female animals. Significant effects are highlighted in dark grey and bolded, trending effects are highlighted in light grey.

| Dependent Variable | Comparison | $F_{(1,57)}$ | p | Effect Size ($\eta_p^2$) | Post-hoc |
|---|---|---|---|---|---|
| PL Bouton Count | Rearing | 1.488 | 0.2276 | 0.025434682 | - |
| | Sex | 0.9889 | 0.3242 | 0.017053139 | - |
| | Treatment | 0.2459 | 0.6219 | 0.004295022 | - |
| | Rearing x Sex | 3.245 | 0.0770 | 0.053856067 | |
| | Rearing x Treatment | 0.005831 | 0.9394 | 0.000102293 | - |
| | Sex x Treatment | 0.7633 | 0.3860 | 0.01321346 | - |
| | Rearing x Sex x Treatment | 2.543e-005 | 0.9960 | 4.46186E-07 | - |
| PL Bouton Intensity | **Rearing** | **10.87** | **0.0017** | 0.160201338 | - |
| | Sex | 0.4126 | 0.5232 | 0.007187106 | - |
| | Treatment | 1.644 | 0.2050 | 0.028032461 | - |
| | **Rearing x Sex** | **15.00** | **0.0003** | 0.208293414 | **CON Male CNO vs. MS Male CNO P=0.0008** |
| | Rearing x Treatment | 2.177 | 0.1456 | 0.036792843 | - |
| | Sex x Treatment | 0.9066 | 0.3450 | 0.015655768 | - |
| | Rearing x Sex x Treatment | 0.003745 | 0.9514 | 6.56889E-0 | - |
| IL Bouton Count | Rearing | 0.2912 | 0.5916 | 0.00508253 | - |
| | Sex | 0.0001787 | 0.9894 | 3.13564E-06 | - |
| | Treatment | 1.631 | 0.2068 | 0.027816011 | - |
| | Rearing x Sex | 3.617 | 0.0623 | 0.059652333 | - |
| | Rearing x Treatment | 0.1810 | 0.6721 | 0.00316502 | - |
| | Sex x Treatment | 1.752 | 0.1909 | 0.029820241 | - |
| | Rearing x Sex x Treatment | 0.9495 | 0.3340 | 0.01638525 | - |
| IL Bouton Intensity | **Rearing** | **15.58** | **0.0002** | 0.214677883 | - |
| | Sex | 0.3025 | 0.5845 | 0.005278463 | - |
| | Treatment | 1.935 | 0.1696 | 0.032831951 | - |
| | **Rearing x Sex** | **13.19** | **0.0006** | 0.187938684 | **CON Male CNO vs. MS Male CNO P=0.0004** |
| | Rearing x Treatment | 1.889 | 0.1747 | 0.032081121 | - |
| | Sex x Treatment | 0.4053 | 0.5269 | 0.007060988 | - |
| | Rearing x Sex x Treatment | 0.004526 | 0.9466 | 7.94012E-05 | - |

Supplementary Table 5. Results of 3-way ANOVAs for sex, rearing, and treatment effect on axonal bouton outcome measures in the prefrontal cortex. Significant effects are highlighted in dark grey and bolded, trending effects are highlighted in light grey.

| Dependent Variable | Comparison | $F_{(1,25)}$ | p | Effect Size ($\eta_p^2$) | Post-hoc |
|---|---|---|---|---|---|
| Male PL Bouton Count | **Rearing** | **4.661** | **0.0407** | 0.157128671 | **-** |
| | Treatment | 0.0957 | 0.3371 | 0.036898273 | - |
| | Interaction | 0.003384 | 0.9541 | 0.000135352 | - |
| Male PL Bouton Intensity | **Rearing** | **36.09** | **<0.0001** | **0.590768089** | - |
| | Treatment | 0.07646 | 0.7844 | 0.003049027 | - |
| | Interaction | 1.404 | 0.2471 | 0.053186702 | - |
| Male IL Bouton Count | **Rearing** | **5.098** | **0.0329** | **0.16938556** | - |
| | Treatment | 0.001856 | 0.9660 | 7.42221E-05 | - |
| | Interaction | 0.2579 | 0.6160 | 0.010209301 | - |
| Male IL Bouton Intensity | **Rearing** | **29.53** | **<0.0001** | **0.541553857** | |
| | Treatment | 0.2925 | 0.5934 | 0.0115662 | |
| | Interaction | 1.6069 | 0.3112 | 0.040991848 | |

Supplementary Table 6. Results of 2-way ANOVAs for rearing and treatment effect on axonal bouton outcome measures in the prefrontal cortex of male animals. Significant effects are highlighted in dark grey and bolded.

| Dependent Variable | Comparison | $F_{(1,33)}$ | p | Effect Size ($\eta_p^2$) | Post-hoc |
|---|---|---|---|---|---|
| Female PL Bouton Count | Rearing | 0.1712 | 0.6818 | 0.005322015 | - |
| | Treatment | 0.07225 | 0.7898 | 0.002252631 | - |
| | Interaction | 0.002575 | 0.9598 | 8.0457E-05 | - |
| Female PL Bouton Intensity | Rearing | 0.1418 | 0.7090 | 0.00441223 | - |
| | Treatment | 2.141 | 0.1532 | 0.06271261 | - |
| | Interaction | 1.013 | 0.3218 | 0.030681587 | - |
| Female IL Bouton Count | Rearing | 0.7413 | 0.3956 | 0.02264177 | - |
| | Treatment | 2.703 | 0.1099 | 0.077893948 | - |
| | Interaction | 0.7832 | 0.3828 | 0.023889501 | - |
| Female IL Bouton Intensity | Rearing | 0.05011 | 0.8243 | 0.001563533 | - |
| | Treatment | 2.072 | 0.1597 | 0.060818423 | - |
| | Interaction | 0.8612 | 0.3603 | 0.026208661 | - |

Supplementary Table 7. Results of 2-way ANOVAs for rearing and treatment effect on axonal bouton outcome measures in the prefrontal cortex of female animals.

| Dependent Variable | Comparison | $F_{(1,79)}$ | p | Effect Size ($\eta_p^2$) | Post-hoc |
|---|---|---|---|---|---|
| OFT- Distance Moved | Rearing | 1.439 | 0.2338 | 0.01789309 | - |
| | Sex | 0.1625 | 0.6880 | 0.0020523 | - |
| | Treatment | 0.9856 | 0.3238 | 0.01232243 | - |
| | Rearing x Sex | 1.514 | 0.2221 | 0.01880951 | |
| | Rearing x Treatment | 0.5380 | 0.4655 | 0.00676355 | - |
| | Sex x Treatment | 3.061 | 0.0841 | 0.03729705 | - |
| | Rearing x Sex x Treatment | .2113 | 0.6470 | 0.00266721 | - |
| OFT-Center Frequency | **Rearing** | **5.353** | **0.0234** | **0.06500756** | **-** |
| | Sex | 0.1538 | 0.6960 | 0.00199353 | - |
| | Treatment | 0.1149 | 0.7356 | 0.00149 | - |
| | Rearing x Sex | 3.594 | 0.0617 | 0.04459166 | |
| | **Rearing x Treatment** | **6.176** | **0.0151** | **0.07423933** | **CON Female SAL vs. MS Female SAL p=0.0266** |
| | Sex x Treatment | 1.611 | 0.2082 | 0.02048872 | - |
| | Rearing x Sex x Treatment | 0.2370 | 0.6278 | 0.00306767 | - |
| OFT-Center Duration | Rearing | 1.587 | 0.2115 | 0.0201978 | - |
| | Sex | 0.6009 | 0.4406 | 0.00774442 | - |
| | Treatment | 1.654 | 0.2023 | 0.02102213 | - |
| | Rearing x Sex | 2.022 | 0.1590 | 0.02559105 | - |
| | **Rearing x Treatment** | **4.912** | **0.0296** | **0.0599623** | **-** |
| | Sex x Treatment | 1.018 | 0.3162 | 0.01304798 | - |
| | Rearing x Sex x Treatment | 0.2226 | 0.6384 | 0.0028821 | - |
| OFT-Latency to Center | Rearing | 2.511 | 0.1171 | 0.0315859 | - |
| | **Sex** | **7.019** | **0.0098** | **0.08353889** | **-** |
| | **Treatment** | **7.148** | **0.0092** | **0.08494022** | **-** |
| | Rearing x Sex | 0.1024 | 0.7498 | 0.00132815 | - |
| | **Rearing x Treatment** | **5.098** | **0.0268** | **0.06209889** | **CON Female SAL vs. CON Female CNO p=0.0206** |
| | Sex x Treatment | 0.7030 | 0.4044 | 0.00904446 | - |
| | Rearing x Sex x Treatment | 0.2234 | 0.6378 | 0.00289258 | - |

Supplementary Table 8. Results of 3-way ANOVAs for sex, rearing, and treatment effect on behavioral outcome measures in the open field test (OFT). Significant effects are highlighted in dark grey and bolded; trending effects are highlighted in light grey.

| Dependent Variable | Comparison | $F_{(1,38)}$ | p | Effect Size ($\eta_p^2$) | Post-hoc |
|---|---|---|---|---|---|
| Male OFT- Distance Moved | Rearing | 0.0004016 | 0.9841 | 0.01549859 | - |
| | Treatment | 0.2406 | 0.6266 | 1.0568E-05 | - |
| | Interaction | 0.5982 | 0.4440 | 0.00629224 | - |
| Male OFT- Center Frequency | Rearing | 0.07037 | 0.7923 | 0.0018985 | - |
| | Treatment | 0.3488 | 0.5584 | 0.00933868 | - |
| | Interaction | 3.561 | 0.0670 | 0.0877695 | - |
| Male OFT- Center Duration | Rearing | 0.01107 | 0.9168 | 0.00029108 | - |
| | Treatment | 2.216 | 0.1448 | 0.05510296 | - |
| | Interaction | 1.281 | 0.2649 | 0.03260555 | - |
| Male OFT- Latency to Enter the Center | Rearing | 2.125 | 0.1536 | 0.05573258 | - |
| | **Treatment** | **4.475** | **0.0414** | **0.1105448** | **-** |
| | **Interaction** | **4.235** | **0.0469** | **0.10524967** | **CON SAL vs CON CNO p=0.0129 CON SAL vs MS SAL p=0.0258** |

Supplementary Table 9. Results of 2-way ANOVAs for rearing and treatment effects on behavioral outcome measures in the open field test (OFT) for male rats. Significant effects are highlighted in dark grey and bolded; trending effects are highlighted in light grey.

| Dependent Variable | Comparison | $F_{(1,41)}$ | p | Effect Size ($\eta_p^2$) | Post-hoc |
|---|---|---|---|---|---|
| Female OFT- Distance Moved | Rearing | 3.555 | 0.0665 | 0.07979239 | - |
| | **Treatment** | **4.526** | **0.0394** | **0.09942083** | **-** |
| | Interaction | 0.04513 | 0.8328 | 0.00109943 | - |
| Female OFT- Center Frequency | **Rearing** | **11.24** | **0.0018** | **0.21930326** | **-** |
| | Treatment | 1.640 | 0.2077 | 0.03938723 | - |
| | Interaction | 2.533 | 0.1194 | 0.05954388 | - |
| Female OFT- Center Duration | **Rearing** | **4.392** | **0.0427** | **0.10119522** | **-** |
| | Treatment | 0.04685 | 0.8298 | 0.0011998 | - |
| | **Interaction** | **4.412** | **0.0422** | **0.10162472** | **CON SAL vs MS SAL p=0.0045** |
| Female OFT- Center Latency to Enter | Rearing | 1.231 | 0.2737 | 0.02915066 | - |
| | **Treatment** | **4.185** | **0.0472** | **0.09262637** | **-** |
| | Interaction | 2.530 | 0.1194 | 0.05812613 | - |

Supplementary Table 10. Results of 2-way ANOVAs for rearing and treatment effects on behavioral outcome measures in the open field test (OFT) for female rats. Significant effects are highlighted in dark grey and bolded; trending effects are highlighted in light grey.

| Dependent Variable | Comparison | $F_{(1,76)}$ | p | Effect Size ($\eta_p^2$) | Post-hoc |
|---|---|---|---|---|---|
| Latency to Startle-Day 1 Baseline | Rearing | 0.8468 | 0.3604 | 0.01101861 | - |
| | **Sex** | **7.821** | **0.0065** | **0.09331452** | **-** |
| | Treatment | 0.1455 | 0.7040 | 0.00191009 | - |
| | Rearing x Sex | 2.425 | 0.1236 | 0.03092006 | |
| | Rearing x Treatment | 0.07213 | 0.7890 | 0.00094787 | - |
| | Sex x Treatment | 0.5432 | 0.4634 | 0.00709585 | - |
| | Rearing x Sex x Treatment | 0.7483 | 0.3897 | 0.00974839 | - |
| Peak Time-Day 1 Baseline | Rearing | 0.8638 | 0.3556 | 0.01123722 | - |
| | Sex | 0.8543 | 0.3583 | 0.01111674 | - |
| | Treatment | 0.02980 | 0.8634 | 0.00039198 | - |
| | Rearing x Sex | 1.638 | 0.2045 | 0.02109971 | |
| | Rearing x Treatment | 0.4685 | 0.4957 | 0.00612809 | - |
| | Sex x Treatment | 0.5485 | 0.4612 | 0.00716628 | - |
| | Rearing x Sex x Treatment | 1.485 | 0.2268 | 0.01916122 | - |
| Peak Value-Day 1 Baseline | Rearing | 0.1371 | 0.7122 | 0.00180069 | - |
| | **Sex** | **5.598** | **0.0205** | **0.0686076** | **-** |
| | Treatment | 1.409 | 0.2390 | 0.0181961 | - |
| | Rearing x Sex | 0.1726 | 0.6790 | 0.00226561 | |
| | Rearing x Treatment | 0.2504 | 0.6182 | 0.00328425 | - |
| | Sex x Treatment | 0.09268 | 0.7616 | 0.00121796 | - |
| | Rearing x Sex x Treatment | 0.01196 | 0.9132 | 0.00015741 | - |

Supplementary Table 11. Results of 3-way ANOVAs for sex, rearing, and treatment effect on acoustic startle outcome measures for baseline performance on Day 1 of testing. Significant effects are highlighted in dark grey and bolded.

| Dependent Variable | Comparison | $F_{(1,76)}$ | p | Effect Size ($\eta_p^2$) | Post-hoc |
| --- | --- | --- | --- | --- | --- |
| Latency to Startle-Day 2 | **Rearing** | **4.855** | **0.0306** | **0.06004806** | **-** |
| | **Sex** | **4.082** | **0.0469** | **0.05097292** | **-** |
| | Treatment | 0.1870 | 0.6667 | 0.00245386 | - |
| | Rearing x Sex | 0.1141 | 0.7364 | 0.00149899 | |
| | Rearing x Treatment | 0.01342 | 0.9081 | 0.00017657 | - |
| | Sex x Treatment | 0.05270 | 0.8191 | 0.00069288 | - |
| | Rearing x Sex x Treatment | 0.08132 | 0.7763 | 0.00106877 | - |
| Peak Time-Day 2 | **Rearing** | **5.210** | **0.0252** | **0.0641583** | **-** |
| | Sex | 0.03596 | 0.5505 | 0.00470834 | **-** |
| | Treatment | 0.2944 | 0.5890 | 0.00385717 | - |
| | Rearing x Sex | 1.061 | 0.3062 | 0.01376994 | |
| | Rearing x Treatment | 0.1908 | 0.6635 | 0.00250387 | - |
| | Sex x Treatment | 0.08678 | 0.7691 | 0.00114037 | - |
| | Rearing x Sex x Treatment | 0.4668 | 0.4965 | 0.00610453 | - |
| Peak Value-Day 2 | Rearing | 0.05308 | 0.8184 | 0.00069794 | - |
| | **Sex** | **10.54** | **0.0017** | **0.12181925** | **-** |
| | Treatment | 0.6101 | 0.4372 | 0.00796385 | - |
| | Rearing x Sex | 1.245 | 0.2680 | 0.01612045 | |
| | Rearing x Treatment | 1.274 | 0.2625 | 0.01648828 | - |
| | Sex x Treatment | 0.04553 | 0.8316 | 0.0005987 | - |
| | Rearing x Sex x Treatment | 0.1949 | 0.6601 | 0.00255811 | - |

Supplementary Table 12. Results of 3-way ANOVAs for sex, rearing, and treatment effect on acoustic startle outcome measures for Day 2 performance following USV presentation. Significant effects are highlighted in dark grey and bolded.

| Dependent Variable | Comparison | $F_{(1,76)}$ | p | Effect Size ($\eta_p^2$) | Post-hoc |
|---|---|---|---|---|---|
| Latency to Startle-% Change | **Rearing** | **4.081** | **0.0469** | **0.05097198** | **-** |
| | Sex | 0.3714 | 0.5441 | 0.00486421 | - |
| | Treatment | 0.3367 | 0.5635 | 0.00440946 | - |
| | Rearing x Sex | 1.246 | 0.2678 | 0.01612936 | |
| | Rearing x Treatment | 0.001550 | 0.9687 | 2.03908E-05 | - |
| | Sex x Treatment | 0.4108 | 0.5235 | 0.005376161 | - |
| | Rearing x Sex x Treatment | 0.2598 | 0.6117 | 0.003406524 | - |
| Peak Time-% Change | **Rearing** | **5.195** | **0.0255** | **0.063984623** | **-** |
| | Sex | 0.9407 | 0.3352 | 0.012226299 | - |
| | Treatment | 0.1547 | 0.6952 | 0.002031252 | - |
| | Rearing x Sex | 2.756 | 0.1010 | 0.034995887 | |
| | Rearing x Treatment | 0.05232 | 0.8197 | 0.00068789 | - |
| | Sex x Treatment | 0.08566 | 0.7706 | 0.001125814 | - |
| | Rearing x Sex x Treatment | 0.2905 | 0.5915 | 0.003808249 | - |
| Peak Value-% Change | Rearing | 0.4270 | 0.5154 | 0.00558611 | - |
| | **Sex** | **4.856** | **0.0306** | **0.060060504** | **-** |
| | Treatment | 1.597 | 0.2102 | 0.020577619 | - |
| | Rearing x Sex | 0.2291 | 0.6336 | 0.003005388 | |
| | Rearing x Treatment | 0.3825 | 0.5382 | 0.005007741 | - |
| | Sex x Treatment | 0.01731 | 0.8957 | 0.000227687 | - |
| | Rearing x Sex x Treatment | 0.02460 | 0.8758 | 0.000323593 | - |

Supplementary Table 13. Results of 3-way ANOVAs for sex, rearing, and treatment effect on percent change of acoustic startle outcome measures from Day 1 to Day 2. Significant effects are highlighted in dark grey and bolded.

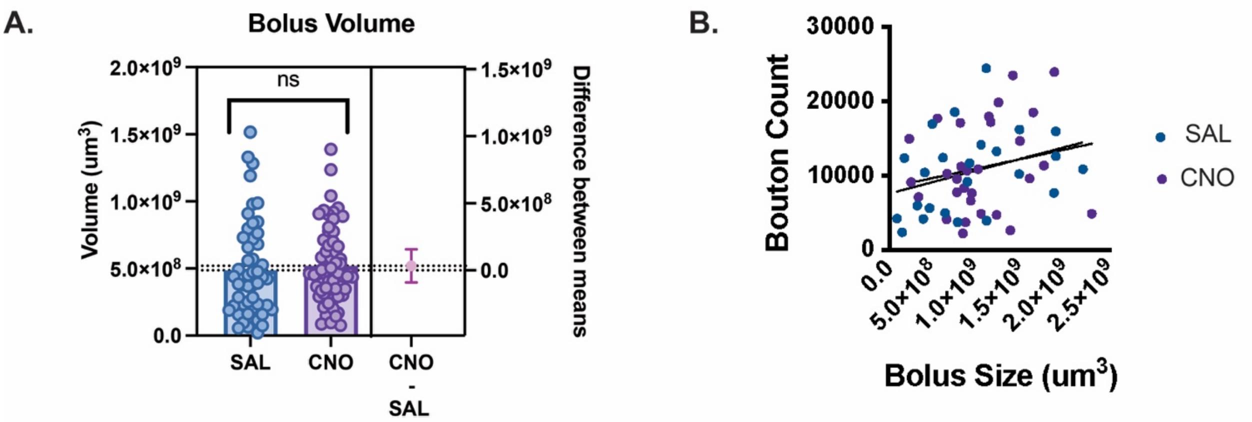


**Supplemental Figure 1.** Additional DREADD validation. (**A**) Bolus volumes do not differ between saline and CNO treated animals ($p>0.05$). n=50-59. (**B**) Bolus size does not correlate with prefrontal cortex bouton count ($p>0.05$) n=25-29.
